# The Anaerobic Efflux Pump MdtEF-TolC Confers Resistance to Cationic Biocides

**DOI:** 10.1101/570408

**Authors:** Diego Novoa, Otakuye Conroy-Ben

## Abstract

The *E. coli* RND transporter MdtEF-TolC is a tri-partite efflux pump that exports toxic substances. Little is known of the full range of substrate specificity of the anaerobic efflux pump, but MdtF shares similar homology and substrate specificity to the major RND efflux protein AcrB. To determine the substrate range of the anaerobic efflux pump MdtEF-TolC, *E. coli* mutants were exposed to 210 different biocides, and growth was monitored. This approach was used to validate AcrAB-TolC substrates and discover new chemicals transported by the major antibiotic efflux protein. Results showed that overexpression of MdtEF conferred resistance to the same substrates as AcrAB-TolC, but were limited to cationic amino-based biocides. Alignment of the amino acids lining the distal pocket of MdtF and AcrB revealed a more acidic isoelectric point (pI) of an order of magnitude in MdtF, whereas the proximal pocket and external cleft were homologous and displayed identical pIs. This analysis suggests that pH, which determines acid-base speciation, and the distal pocket surface proteins play a role in MdtF substrate specificity.

**Importance:** High-throughput screening of *E. coli* mutants revealed that the substrates of the anaerobic efflux pump MdtEF-TolC are the same cationic biocides exported by AcrAB-TolC. Comparison of the protein sequences of the distal pocket, proximal pocket, and external cleft of the two RND proteins showed homology in amino acid surface charge and isoelectric point. Residue differences within the distal pocket are responsible for a more acidic pI and greater negative charge of the inner membrane protein MdtF surface, and support the findings of transport of cationic substances.

## Introduction

*E. coli* expresses multi-drug efflux proteins in response to intracellular toxin accumulation. These pumps are of the Resistance Nodulation Division (RND), ATP-Binding-cassette (ABC-type), multi-drug and toxic compound extrusion (MATE) and small multi-drug resistance (SMR) types. The major multi-drug efflux protein, AcrAB-TolC, is an RND pump responsible for extrusion of a number of chemicals, including bile salts, hormones, and antibiotics. The other six RND efflux pumps (CusCFBA, AcrEF, AcrD, MdtEF, and YegMNO) also work to export monovalent cations (CusCFBA), copper and zinc (AcrD), aminoglycosides (AcrD), and those similar to AcrAB (YhiUV and AcrEF) (1–6). The substrates of MdtEF (also known as YhiUV)-TolC are limited in scope, but are similar to the major multidrug efflux pumps AcrAB-TolC of *E. coli* and MexAB-OprM in *Pseudomonas aeruginosa*. They include the steroid hormones, deoxycholate, cationic detergents, crystal violet, ethidium, and erythromycin, reviewed in (7). A protein blast of MdtF (Accession P37637) the inner membrane protein of the RND complex responsible for substrate recognition and binding, is most homologous to AcrB (71% identity) and MexB (63% identity, unpublished, our work). The protein identity is indicative of similar substrates across protein classes and bacteria species.

While AcrAB-TolC is the major multidrug efflux pump that is constitutively expressed in *E. coli*, the role of YhiUV-TolC is complex. GadX, a regulator of acid resistance, can induce MdtF expression through GadE (8, 9) (10, 11). MdtEF is also regulated by ArcA under anerobic growth (8). MdtEF is expressed differentially during lag and stationary growth, and under toxic stress (12). Our previous work showed that the environmental pollutants 17α-ethynylestradiol, bisphenol A and nonylphenol induce expression of both AcrB and MdtF (13). The pump has been termed the anaerobic efflux pump due to increased expression and efflux in anaerobic conditions (8, 9, 14).

In this work, we determined the substrates of MdtF and AcrB using a high-throughput antimicrobial screen. *E. coli* mutants (Table 1) were exposed to 210 chemicals found in Biolog’s Chemical Sensitivity Panels covering different biocide classes was used in the growth comparison. Additional chemical data (pKA, molecular charge) were extracted from Chemicalize for substrates of interest to both proteins. An alignment of the inner membrane proteins AcrB and MdtF provided insight into the surface chemistry of the binding pockets of the RND proteins.

**Table 1:** Chemical classes found in Biolog’s chemical sensitivity panels.

| Location of action | Antimicrobial class |
| --- | --- |
| Chelator | Carboxylic acids, hydroxyquinolines, N-heterocycles |
| DNA & RNA | Alkylation, fluoroquinolones, intercalators, nitrofurans analogs, purine/pyrimidine analogs, quinolones |
| Folate | Sulfonamides |
| Fungicide | Lipoxygenases, phenylsulfamides |
| Ion channel | K<sup>+</sup> inhibitors, Na<sup>+</sup> inhibitors |
| Membrane | Anionic/cationic/zwitterionic detergents, electron transport, guanidine (permeability), phenothiazines (efflux pump inhibitor), |
| Other biocide | Anti-capsules, acetylcholine antagonist, fatty acid synthesis, glycopeptides, nitroimidazoles, rifamycins, triazoles |
| Oxidation | Glutathiones, oxidizing agents, sulfhydryl |
| Protein | Aminoglycosides, lincosamides, macrolides, tetracyclines, tRNA synthetase |
| Respiration | Ca<sup>2+</sup> transporters, ionophores, uncouplers |
| Wall | Cephalosporins, monobactams, β-lactams, peptidoglycan synthesis, polymyxins, glycopeptides |

## Results

### Overexpressed AcrAB

The classes of chemicals transported by AcrAB-TolC are shown in Table 2 and many have been reported elsewhere (3, 7, 15). However, with this screen, we verified additional chemicals within the macrolides, tetracyclines, cationic membrane detergents, chloramphenicol and thiamphenicol, fluoroquinolones, and quinolones. New chemical and classes uncovered by this assay were the metal-chelating 1,10-phenanthroline, unsubstituted and halogenated hydroxyquinolines, phenothiazines, glycopeptides, triclosan, and pentachlorophenol among others. With these reproducible results, and for those comparable to literature, interpretation of two-fold growth is an appropriate metric for evaluating antibiotic susceptibility.

**Table 2:** Substrates of AcrAB and MdtEF determined from two-fold growth differences of *E. coli* mutants, with charges on molecule extracted from Chemicalize. **DNA intercalators acriflavine, proflavine, and 9-aminoacridine are likely substrates of MdtEF-TolC, however different hosts are needed to validate this.

| Class | (Charge): Chemicals | AcrB | MdtF |
| --- | --- | --- | --- |
| Cationic detergent | (+1): benzethonium chloride, cetylpyridinium chloride, dequalinium chloride, dodecyltrimethyl ammonium bromide, domiphen bromide, methyltrioctylammonium chloride | X | X |
| Macrolide | (+1): josamycin, oleandomycin, spiramycin, troleandomycin, tylosin, erythromycin | X | X |
| Other biocides | (-1): niaproof; (0): 2,4-diamino-6,7-diisopropyl-pteridine (0/129); (+1): lidocaine, dodine, amitriptyline, pridinol, sanguinarine, chelerythrine, orphenadrine, trimethoprim | X | X |
|  | (-1): hexachlorophene, rifamycin SV, lawsone, fusidic acid, pentachlorophenol; (0): nordihydroguaiaretic acid, lauryl sulfobetaine, triclosan, harmene, lincomycin | X |  |
| Phenothiazine | (+1): thioridazine, trifluoperazine, chlorpromazine | X | X |
| Glycopeptide | (+1): phleomycin | X |  |
|  | (+2): bleomycin | X | X |
| Respiration uncoupler | (+1): crystal violet | X | X |
|  | (0): iodonitrotetrazolium violet, menadione | X |  |
| Chelator | (0): 5-chloro-7-iodo-8-hydroxyquinoline | X | X |
|  | (0): 1,10-Phenanthroline, 5,7-dichloro-8-hydroxyquinaldine, 5,7-dichloro-8-hydroxyquinoline, 8-hydroxyquinoline | X |  |
| DNA intercalator | (-1): novobiocin; (0): 2-phenylphenol; (+1): acriflavine, proflavine; (+2): 9-aminoacridine | X | ** |
| Fenicol | (0): chloramphenicol, thiamphenicol | X |  |
| Fluoroquinolone | (-1): ofloxacin; (0): ciprofloxacin, enoxacin, lomefloxacin, norfloxacin | X |  |
| Quinolone | (-1): nalidixic acid, oxolinic acid | X |  |
| Tetracycline | (0): chlortetracycline, doxycycline, oxytetracycline, rolitetracycline, tetracycline | X |  |

### Overexpressed MdtF

The screen showed that MdtF substrates were the same substrates as AcrB (Table 3), but were limited to cationic biocides. Growth was two-fold greater than control for overexpressed MdtEF for cationic detergents, macrolides, the metal chelators clioquinol and sanguinarine, 2,4-diamino-6,7-diisopropyl-pteridine, amitryptiline, the phenothiazines (efflux pump inhibitors), pridinol, chelerythrine, crystal violet, and cefmetazole. There were no differences in substrate specificity at t = 16 hours for *E. coli* grown aerobically or anaerobically growth, except that anaerobically cultures grew ^slower. The slower growth was accounted for by determining relative growth, where OD600 was^ normalized to bacteria grown with no antibiotics.

**Figure 1:**
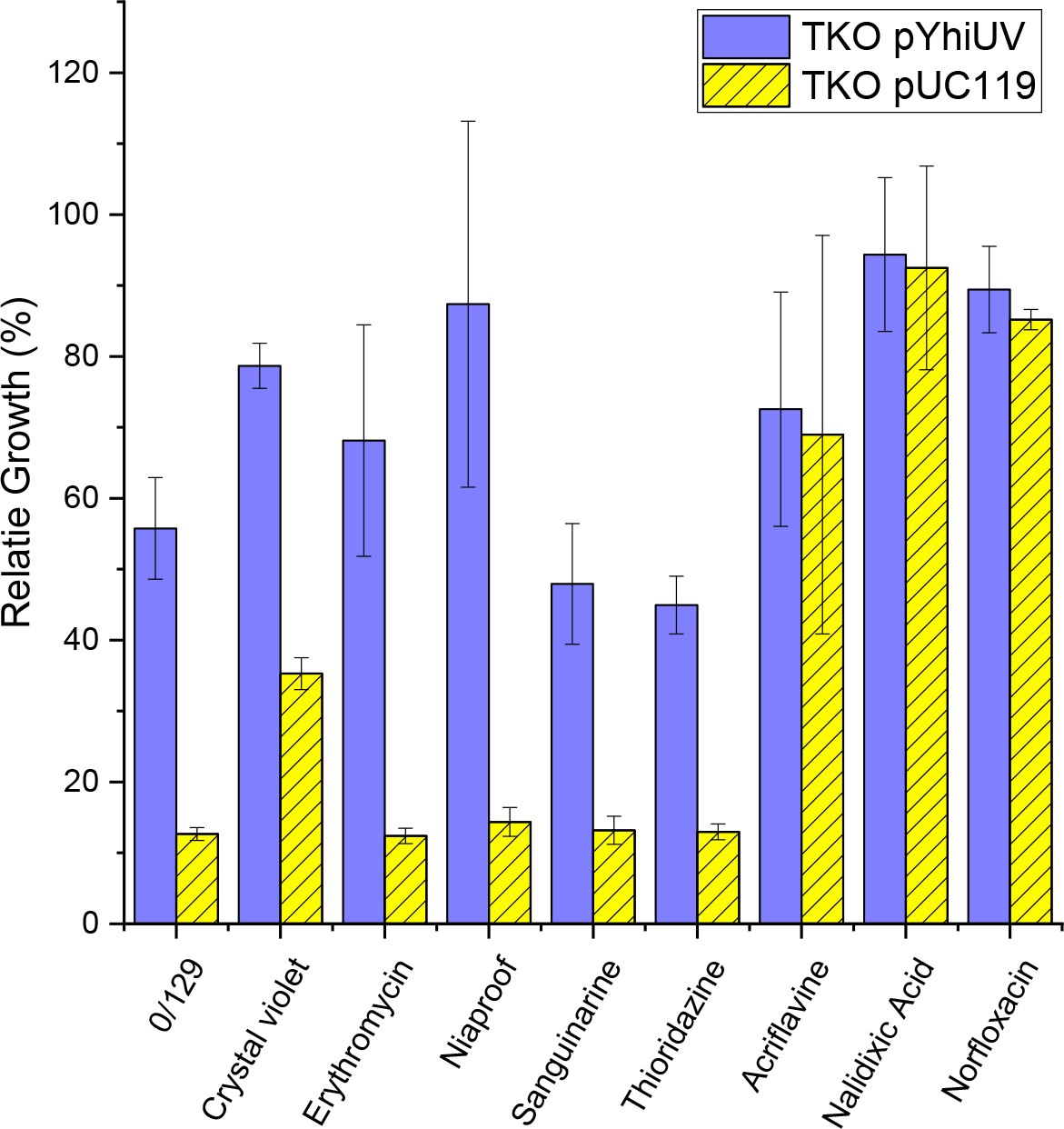
Relative growth of TKO pMdtEF and TKO pUC119 on select antibiotics from Biolog chemical sensitivity panels. Antibiotics were considered substrates if the gene insert (dark blue bars) allowed for two-fold greater growth than that by the empty vector (yellow bars). Acriflavine, nalidixic acid, and norfloxacin were not considered substrates, nor are they reported in literature. Error bars represent standard deviation of three independent experiments.

**Table 3:**
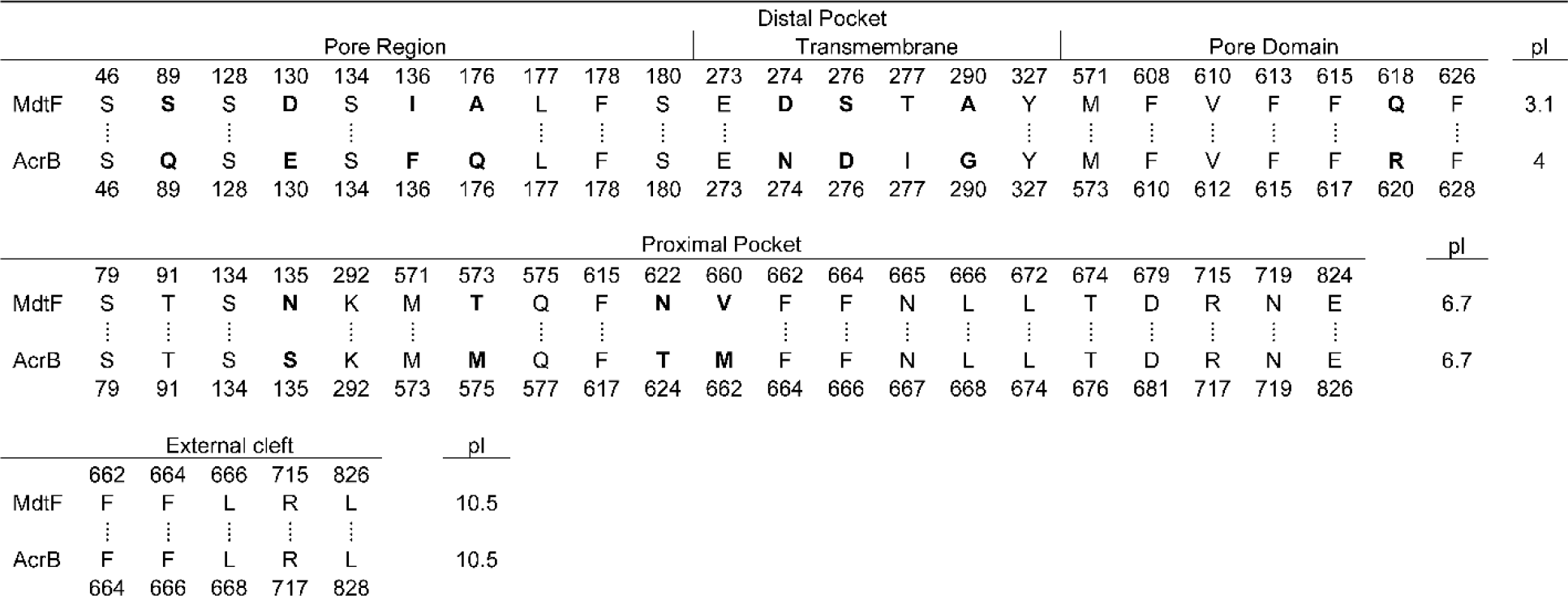
Binding domain alignment. Critical residues extracted from AcrB (described in (20)), with corresponding positions, are homologous to MdtF. Differences within the distal pocket are responsible for a more acidic pI and greater negative charge of MdtF.

Closer analysis of the chemicals that were substrates of the pump MdtEF and not AcrAB showed a trend according to structure and chemical property. We looked further at properties of the AcrAB substrates using the Chemicalize computational database (16, 17), focusing on the pKa, structure, and charge at test pH. We found that the substrates of MdtEF are also the substrates of AcrAB that have a +1 charge (Figure 2). A statistical analysis of the charge of MdtEF versus AcrAB substrates showed that the anaerobic efflux pump exports primarily cations, while AcrAB has broad substrate specificity. There were exceptions to the +1 charge rule (n = 2), however the average charge for AcrAB chemicals and that for MdtEF were statistically significant (p = 0.00065).

**Figure 2:**
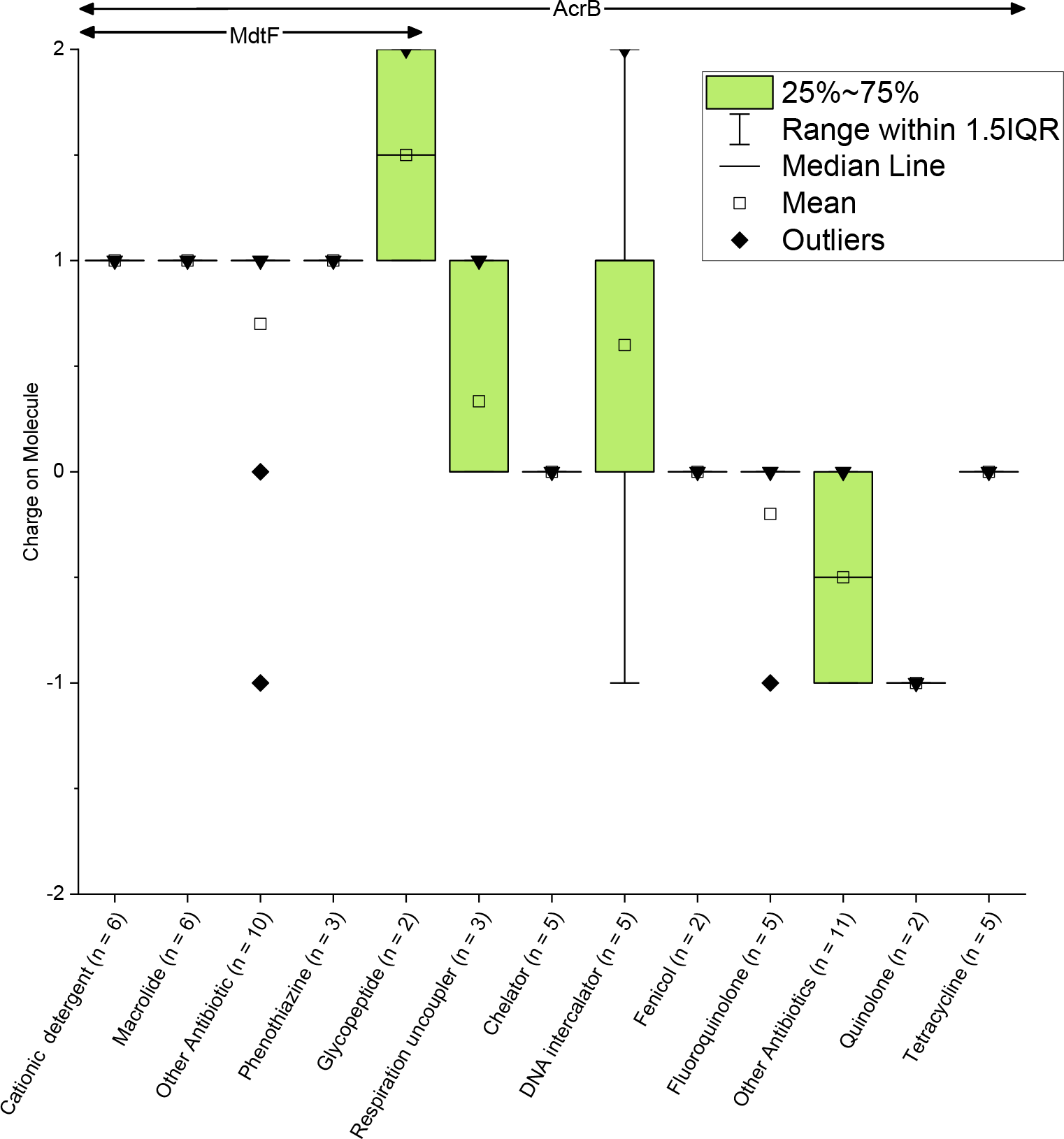
Charges of biocide classes exported by MdtEF and AcrAB. The two outlying non-cationic points under “Other antibiotic” that are MdtEF substrates are Niaproof, an anionic surfactant, and 2,4-diamino-6,7-diisopropyl-pteridine. IQR = interquartile range.

## Discussion

Due to substrate similarity and known homology between various RND inner membrane proteins, the sequences of MdtF (accession P37637) and AcrB (accession P31224) were compared (18). Protein alignment showed 71% identity, 84% positives, and two gaps (Supplemental file S1). The periplasmic domain (residues 33-335) alignment also showed homology, with 67% identity, 83% positive, and no gaps. The isoelectric points (19) within this region were the same at 4.4. A second periplasmic region (residues 565-871 of AcrB) showed differences between the two proteins, but the overall isoelectric points (pI) for MdtF and AcrB are 4.9 and 4.8 with 65% identity and 79% positives. Generally, the proteins exhibit similar chemistries, however, there are specific regions within the bulk protein that require further analysis based on *in silico* studies of AcrB channels (20).

Looking closer at the distal pocket, proximal pocket, and external cleft, identified as essential binding regions in AcrB (20), surface residues were aligned with MdtF, which displayed similar homology overall (Table 3). However, slight differences in the distal pocket suggest the local chemistry may play a role in substrate specificity. MdtF residues lining the distal pocket were more acidic (pI = 3.1) than AcrB (pI = 4.0). Further, AcrB residues were more diverse, covering polar, aliphatic, neutral, aromatic, and positive amino acid classes, while the MdtF surface was restricted to polar, aliphatic, and neutral residues. At pH = 7.4, the charge of MdtF surface residues was -3, while the charge of AcrB was -2. It has been reported that the distal pocket is responsible for substrate specificity and direct interactions between protein and substrate, and our results show that may be so in MdtF as well. With respect to other essential binding regions, differences were observed in the proximal pocket, however the amino acid classes were maintained over the four residues, with no change in the isoelectric point. The external cleft surface sequence and pI between AcrB and YhiV were the identical.

The substrates exported by AcrB are diverse in structure, size, charge, and hydrophobicity, while the substrates of MdtEF are limited to cations along with a few exceptions. Reported exceptions include neutral and anionic substrates of MdtEF such as bile salts, steroid hormones and hormone mimics, and anionic detergents (13, 15). In this work, two neutral/anionic species were substrates of MdtEF-TolC: Niaproof (anion) and 2,4-Diamino-6,7-diisopropyl-pteridine (neutral). Niaproof is an anionic detergent similar in structure to sodium dodecyl sulfate (SDS), also transported by MdtEF. 2,4-Diamino-6,7-diisopropyl-pteridine (neutral) possesses two basic amino functional groups attached to the pteridine ring that were not assigned pKa values by Chemicalize. These amino moieties likely are charged with pKa values lying in physiological pH, but the structure would need to be fully analyzed with a more robust *in silico* approach.

Further, not all cationic substrates of AcrAB were substrates of MdtEF with this screen. In fact, four of the listed cationic substrates of AcrAB were expected to be substrates of MdtF: the similarly structured acriflavine, 9-aminoacridine and proflavine, and the glycopeptide phleomycin. The former are antiseptics that are substrates of at least five other membrane-bound transporters in *E. coli* (2, 21-23), and to truly study conferred resistance in a live host, multiple gene deletions are likely needed. Phleomycin on the otherhand, is a polyprotic glycopeptide with a +1/+2 charge at neutral pH values, and is predominantly neutral at pH = 7.7. An *E. coli* efflux pump does exist for glycopeptides (AmpG) (24), and perhaps this protein more efficient at phleomycin removal than the presence of MdtF.

Our analysis suggests that pH, which determines acid-base speciation and surface charge of the distal pocket, plays a role in MdtF substrate specificity. Isoelectric point analyses showed that the MdtF surface charge is more negative than AcrB. This would allow stronger electrostatic interactions to occur between the channel and cationic substrates. The pKa also determines whether the analyte is charged at a given pH. Twenty-five out of 27 (93%) of the confirmed substrates of MdtEF were cations. However, the two anion/neutral biocides, along with other reported human hormones (neutral) and bile salts (carboxylic and sulfonic acids) are exported. This may be due to a redundancy in substrate specificity since hormones and bile salts are known substrates of the AcrAB-TolC system (2, 25, 26). Bile salts cause widespread protein unfolding and disulfide stress in *E. coli* (27). MdtEF might export such compounds as a response to this added stress. The MdtEF structure may also be responsible for waste metabolite exportation (28). MdtEF may recognizes metabolites from carbon metabolism which may be why some neutral compounds are exported as well. Further studies are needed to explore these variations in substrate specificity.

In this work, we used a high-throughput method to determine MdtEF substrates, which were primarily cationic biocides derived from AcrAB-TolC analytes. Using two-fold growth as a metric for assessing sensitivity, we validated previously published substrates, and discovered new substrates. Collectively, we were able to demonstrate that the presence MdtEF-TolC renders *E. coli* less sensitive to cationic biocides, suggesting the role of MdtEF-TolC as a proton/cation antiporter. The significance of these findings are that under acidic conditions, the lower pI of essential binding residues in the distal pocket would deprotonate, allowing for stronger interactions between cation and anion. These findings suggest that acidic conditions influence the transport of cationic substrates for the exporter MdtEF.

## Materials and Methods

Strains and plasmid descriptions and sources are found in Table 4. Electrocompetent *E. coli* W4680 (kan^R^) and TKO (kan^R^) were transformed with plasmids and plated on 100 mg/L ampicillin in LB agar. After selection, single colonies were pre-cultured in their appropriate antibiotics and inducers (IPTG, 1 mM) and grown to mid-log phase. Plates (Biolog chemical sensitivity panels PM11-20) were seeded with LB media at 5 x 10^5^ cells/mL (200 μL) and grown aerobically or anaerobically at 37 °C. A control plate containing no chemical was also inoculated and grown. Anaerobic conditions were described in (8), where 10 mM nitrate was supplied in LB media as the terminal electron acceptor in LB media. Optical density (λ = 600 nm) was recorded at t = 0 and at t = 16 hours using a platereader (Biotek). Relative growth was calculated as OD_600,antibiotic growth_/OD_600,no chemical growth._

**Table 4:** *E. coli* mutants and plasmids used in this study.

| Strain | Genotype | Source |
| --- | --- | --- |
| W4680A | K-12 ΔacrAB | (29) |
| TKO | K-12 ΔacrAB ΔacrF ΔyhiV | (30) |
| Plasmid |  |  |
| pAB | AcrAB cloned into pUC18, amp<sup>r</sup> | (3) |
| pUC18 | Vector control, amp<sup>r</sup> |  |
| pyhiUV | MdtEF cloned into pUC119, amp<sup>r</sup> | (2) |
| pUC119 | Vector control, amp<sup>r</sup> | (31) |

Chemical data were extracted from the Chemicalize database. pKa was not a property for all biocides, as some chemicals did not possess acidic hydrogens. The pKa (pKa =-log of Ka, the acid dissociation constant) was compared to the test pH (pH = 7.2) to estimate the biocide charge.

### Data Interpretation

Masses present in Biolog’s pre-plated assays are proprietary, however the span of the four concentrations were relevant for studying *E. coli* toxicity, as we observed full growth and no growth over the four concentrations for nearly all analytes, with the exception of tetrazolium violet and puromycin, which were toxic. However, we could not determine the MIC nor the fold-difference concentration from the panel, a method typical of analyzing conferred resistance. Rather, we interpreted results by comparing growth at 16 hours for gene insert and empty vector plasmid. Chemicals were considered substrates of the efflux pump if the growth of the strain with the pump insert was 2-fold greater than the plasmid control for the same chemical concentration in the corresponding well.

